# Paridiprubart inhibits TLR4-dependant NF-κB activation by multiple pathogens

**DOI:** 10.1101/2023.06.27.545921

**Authors:** Ramy Malty, Richard Hilbe, Sang Ahn, Leah Kesselman, Jessica Lam, Karina Kasawara, Larissa Costa, Nishani Rajakulendran, Blair Gordon, Michael Brooks, Samira Mubareka, Ivan Tancevski, Scott D. Gray-Owen

## Abstract

Respiratory pathogens such as SARS-CoV-2 and influenza can activate an exaggerated inflammatory response (cytokine storm) in the lungs that may result in acute respiratory distress syndrome (ARDS), hospitalization, and death. Therapies that target a specific pathogen (i.e. anti-virals) must, by nature, be selected after a specific diagnosis and may become ineffective due to pathogen evolution. An alternate strategy is to counter the exaggerated innate immune response present in ARDS patients using host-directed drug therapies that are agnostic to the infectious agent to overcome both of these challenges. Originally described as the innate immune receptor for lipopolysaccharide (LPS), Toll-like receptor 4 (TLR4) is now understood to be an important mediator of inflammation caused by a variety of pathogen-associated molecular patterns (PAMPs) and host-derived damage-associated molecular patterns (DAMPs). Here we show that paridiprubart, a monoclonal antibody that prevents TLR4 dimer formation, inhibits the response to TLR4 agonists including LPS, the SARS-CoV-2 spike protein, the DAMP high mobility group box 1 (HMGB1), as well as the NF-κB response to infection by both viral and bacterial pathogens. Notable in this regard, we demonstrate that SARS-CoV-2 increases HMGB1 levels, and that paridiprubart inhibits both the SARS-CoV-2 and HMGB1-triggered NF-κB response, illustrating its potential to suppress this self-amplifying inflammatory signal. We also observed that the inhibitory effect of paridiprubart is apparent when cells are exposed to the SARS-CoV-2 spike protein, which is itself a direct TLR4 agonist. In the context of active infection, paridiprubart suppressed the NF-κB-dependent response elicited by infection with SARS-CoV-2, the seasonal coronavirus 229E, influenza A virus or *Haemophilus influenzae*, a gram-negative bacterial pathogen. Combined, these findings reinforce the central role played by TLR4 in the inflammatory response to infection by diverse pathogens, and demonstrates the protective potential of paridiprubart-dependent inhibition of pathogenic TLR4 responses.

## Introduction

Acute Respiratory Distress Syndrome (ARDS) is a complex life-threatening condition of acute respiratory failure due to lung inflammation and oedema [1]. ARDS pathogenesis involves multiple injury, inflammation and coagulation pathways either activated or dysregulated in the lungs and, in some, instances systemically as well [1]. ARDS develops due to either a direct or indirect injury to the alveolar structure of the distal lung and its associated microvasculature. This injury could be due to viral and non-viral infection, sepsis, collagen vascular disease, drug effects, inhalants, shock, acute eosinophilic pneumonia, immunologically mediated pulmonary hemorrhage and vasculitis [2, 3]. This exaggerated immune response has been linked to severe viral pneumonias that lead to pathophysiological organ failure, including avian influenza, seasonal influenza and the severe acute respiratory syndrome (SARS) virus [4-6]. Consistent with this, several studies have demonstrated inflammatory organ injury and ARDS in patients with COVID-19 [7, 8].

Paridiprubart is a monoclonal antibody (mAb) that inhibits signaling by preventing the ligand-induced dimerization of Toll-like receptor 4 (TLR4; [9]) -a key pattern recognition receptor involved in the activation of the innate immune system. Multiple pathogen-associated molecular patterns (PAMPs) and host-derived damage associated molecular patterns (DAMPs) have been discovered to bind and activate TLR4 [10-24], which elicits a pro-inflammatory cytokine response. While this provides an important protective effect, an excessive response can be pathological, leading to a variety of inflammatory conditions, including viral-mediated ARDS [25].

Ligand-induced dimerization of monomeric TLR4 initiates the production of cytokines and chemokines via two separate signaling pathways [26]. The first is the myeloid differentiation factor 88 (MyD88)-dependent pathway, which mediates responses via nuclear factor kappa B (NF-κB) activation, leading to the production of proinflammatory mediators such as tumor necrosis factor alpha (TNF) and interleukins 6 and 8 (IL-6 and IL-8). The second is the Toll/IL-1 receptor (TIR)-domain-containing adaptor-inducing interferon-β (TRIF)-dependent pathway, which promotes the production of type 1 IFNs and the many IFN-dependent genes [27, 28].

There is a growing body of evidence suggesting that viral-mediated cell damage in the lungs leads to the release of DAMPs (e.g., S100A8/A9 (Calprotectin), HMGB1, and oxidized phospholipids (Ox-PL)), which bind to and activate TLR4 [18, 22, 29-31]. Elevated levels of these DAMPs are apparent in the blood of hospitalized patients with COVID-19 and positively correlate with both neutrophil count and disease severity, allowing them to be used as an early predictor of disease severity [29-33]. Moreover, recent studies have shown that the SARS-CoV-2 spike protein can directly bind to TLR4 and stimulate the production of pro-inflammatory cytokines [14, 34, 35]. This effect is not unique to SARS-CoV-2 given that a number of other viral glycoproteins also directly bind and activate TLR4, including those from Ebola virus, Dengue virus, and respiratory syncytial virus (RSV) [10, 11, 36]. When considered together, these findings highlight the potential for TLR4 antagonism as a broadly-effective host-targeted therapeutic strategy with potential utility in both infectious and non-infectious inflammatory disorders.

Here we confirmed [37] that SARS-CoV-2 infection increases levels of the TLR4-activating DAMP HMGB1, and that HMGB1 in turn elicits a pro-inflammatory signaling response. We also show that this increase in TLR4 signaling by SARS-CoV-2 and other pathogens can be inhibited by the anti-TLR4 antibody, paridiprubart. Overall, these findings demonstrate that multiple pathogens can cause an overactive immune response through TLR4 signaling which can be inhibited with paridiprubart treatment.

## Materials and Methods

### Cell culture

THP1-Dual cells (InvivoGen, #thpd-nfis) were grown in RPMI 1640 (Wisent, #350-002-CL) supplemented by 10% FBS (ThermoFisher Scientific, #12483020), 1% Glutamax and β-mercaptoethanol (Sigma Aldrich, #M3148) at a final concentration of 50 mM. Approximately 2×10^6^ frozen cells were recovered in T25 flasks with 5 ml of media and after 3 days cells were counted and diluted to 2-5×10^5^ cells/ml. To differentiate THP-1 dual cells, phorbol 12-myristate 13-acetate (Sigma-Aldrich, # P8139) was added to the complete media at 50 ng/ml final concentration and the cells incubated for 2 hours at 37°C. These cells were then centrifuged at 270 *g* for 5 mins, resuspended in PBS and centrifuged again. Finally, cells were plated at 1×10^5^ cells/well in 96-well plate. Cells are considered differentiated and used after 48 hours.

### SARS-CoV-2 and HCoV-229E propagation

All experiments involving SARS-CoV-2 were conducted in the Toronto High Containment Facility Level 3 lab at the University of Toronto. SARS-CoV-2 was propagated in VeroE6 cells as described before [38]. Briefly, VeroE6 cells were grown to 90% confluence in complete media (high glucose DMEM containing 10% FBS and 1% Pen/Strep). At the time of infection, media was removed, and cells washed with PBS twice. SARS-CoV-2 (isolate SB3, passage 2)

[39] was diluted in DMEM (1% Pen/Strep, without FBS) and this used to replace the PBS to achieve an MOI=0.001. Virus-containing samples were rotated to mix every 15 mins and after 1 hr, this media was removed, and cells were washed with PBS three times. For viral harvesting from a T150 flask, 25 ml of DMEM (1% Pen/Strep, 2% FBS) were added and cells were monitored for the appearance of cytopathic effects, usually within 48-72 hours. When 90% of the cells are dead and floating, the media was harvested and centrifuged at maximum speed for 15 mins to remove cellular debris, and virus-containing supernatant was aliquoted, titrated and stored at -80°C. HCoV-229E is propagated in a similar manner except using Huh7 cells instead of VeroE6 cells and the duration of propagation is usually 24-48 hours.

### Viral (coronaviruses and IAV) and non-typable *Haemophilus influenzae* bacterial infections

For viral infection, cells (differentiated THP1-dual) were washed with PBS and then incubated with serum-free RPMI or DMEM, respectively, containing infectious virus [SARS-CoV-2, MOI 1 [39], HCoV-229E, MOI 1 (ATCC, #VR-740) or IAV MOI 10 (ATCC, #VR-95)] at the desired MOI with gentle rocking every 15 mins. After 1 hr, cells were washed with PBS and then incubated with growth medium containing 2% FBS for 48 hrs.

In the case of non-typeable *Haemophilus influenzae* (NTHi) infection [40], the bacteria was grown to OD_600_ of approximately 0.5 and added directly to THP1-dual cells at 1 CFU/cell for the entire duration of the experiment.

### ELISA and reporter assay

Media was harvested from the infected or mock-treated cell cultures and subjected to centrifugation at maximum speed in 96-well V-bottom plate for 15 mins before being carefully transferred into 96-well plate ELISA plates without contacting the bottom of the well. Human HMGB1/HMG-1 ELISA (Novus Biologicals, NBP2-62766) was performed as described in manufacturer’s instructions. For the secreted alkaline phosphatase-dependent NF-κB reporter assay, 20 µl of media from the THP-1 Dual cells was incubated with 180 µl Quanti-Blue (InvivoGen, # rep-qbs) and incubated for 4 hours at 37°C before quantification at OD_635_. ELISA and reporter assay quantification was done using Synergy H1 plate reader fitted with Gen5 software.

### RNA extraction and quantitative real-time PCR

Total cellular RNA extraction from cells grown in a 24-well plate was performed using RNeasy Plus kit (Qiagen, # 74034) as per manufacturer’s protocol, with RNA finally eluted using nuclease-free water and then quantified using a NanoDrop One (ThermoFisher Scientific).

Reverse transcriptase qPCR was performed using Luna Universal Probe One-Step RT-qPCR Kit (NEB, # E3006L). The following predesigned primers and probe sets for each target and the housekeeping gene GAPDH were purchased from IDT:

HMGBI (Primer 1: 5’-GGATCTCCTTTGCCCATGT-3’, Primer 2: 5’-CAGCCATTGCAGTACATTGAG-3’, Probe: 5’-/56-FAM/ACAGAGTCG/ZEN/CCCAGTGCCC/3IABkFQ/-3’);

GADPH (Primer 1: 5’-GCGCCCAATACGACCAA-3’, Primer 2: 5’-

CTCTCTGCTCCTCCTGTTC-3’, Probe: 5’-

/5SUN/CCGTTGACT/ZEN/CCGACCTTCACCTT/3IABkFQ/-3’).

To enable multiplexing of each target gene with housekeeping gene, probes were FAM-and SUN-labelled, respectively.

### *Haemophilus influenzae* growth curve and infection

Non-typable *Haemophilus influenzae* (NTHi, strain 86028 NP^2^) was grown from frozen stocks on chocolate agar (Hardy Diagnostics, # E14) at 37°C 5% CO^2^. Bacteria were subcultured from the chocolate agar into brain heart infusion (BHI) broth supplemented with 1% BBL IsoVitaleX (BD Biosciences, # 211875) to a starting OD600=0.1. These cultures were shaken at 200 rpm and 37°C to obtain an OD_600_ ≈ 0.3 to ensure bacterial cells are metabolically active and in exponential growth phase when inoculating onto the THP1-dual cells.

## Results

### SARS-CoV-2 infection induces production of DAMPs *in vitro*

Multiple COVID-19 patient cohort studies have shown that serum levels of TLR4-specific DAMPs such as HMGB1 are elevated in hospitalized COVID-19 patients, positively correlate with both neutrophil count and disease severity and, importantly, function as an early predictor of disease progression [29-33]. Given these findings, we first tested the effects of SARS-CoV-2 infection on HMGB1 production *in vitro*. SARS-CoV-2 infection significantly increased transcriptional expression of HMGB1 in the THP1 monocytic cell line (**Figure 1a**). Furthermore, SARS-CoV-2 infection significantly increased HMGB1 protein release by THP1 cells (**Figure 1b**). These findings indicate that SARS-CoV-2 infection can directly increase production of the alarmin, HMGB1.

**Figure 1.**
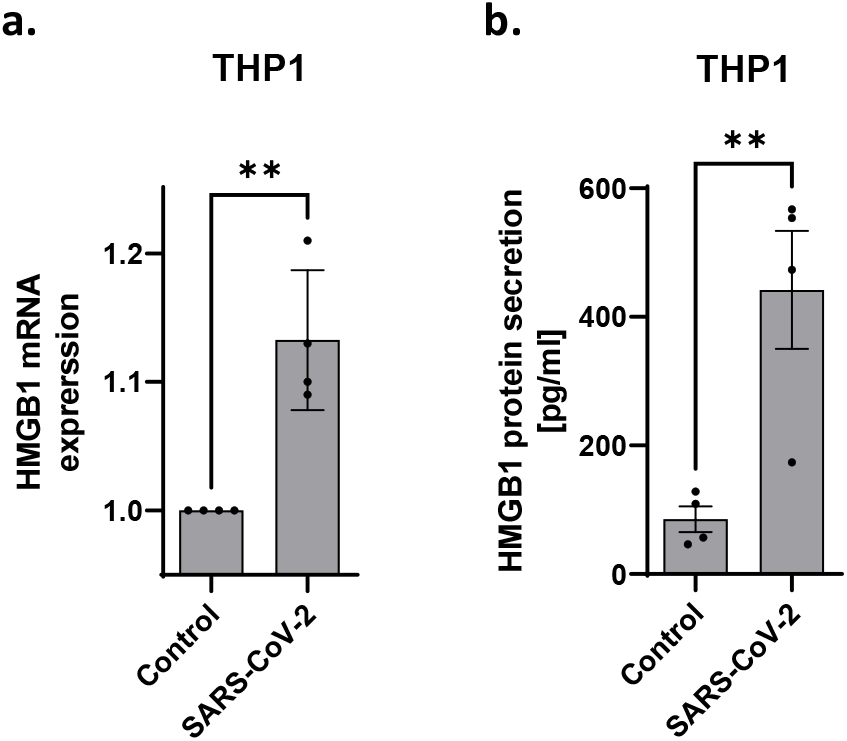
SARS-CoV-2 increases HMGB1 expression in monocytic and epithelial-derived cells. **A)** Infection of PMA-differentiated THP1 monocytes (right panel) with SARS-CoV-2 at MOI 10 results in a significant increase in relative HMGB1 mRNA expression as determined by RT-qPCR. **B)** Infection of PMA-differentiated THP1 cells with SARS-CoV-2 at MOI 10 results in a significant increase in HMGB1 protein release in cell culture media as determined by ELISA. **p<0.01

### Paridiprubart blocks the activation of NF-κB pathway by PAMPs and DAMPs

TLR4 is the most extensively studied member of the TLR family [41]. While originally considered to function primarily as the receptor for bacterial-derived LPS, TLR4 can bind and become activated in response to PAMPs and DAMPs of diverse type and origin. Binding of these agonists to TLR4 triggers receptor dimerization to initiate intracellular signaling that leads to the expression of pro-inflammatory cytokine and interferon expression [22]. Paridiprubart is a humanized immunoglobulin gamma (IgG) 1 kappa (κ) mAb that binds and antagonizes TLR4 in a ligand-independent manner, preventing the ligand-driven receptor dimerization and downstream signaling cascades. Using the TLR4 reporter cell line (THP1-Dual_TM_), which allows NF-κB transcription factor activity to be monitored by measuring secreted embryonic alkaline phosphatase (SEAP) activity in the culture supernatant, we confirmed that activation of TLR4-mediated NF-kB signaling by LPS can be inhibited by paridiprubart in a dose-dependent manner (**Figure 2a**). Previous studies have suggested that SARS-CoV-2 can activate TLR4 signaling via two mechanisms [14, 15, 29, 34, 42]. One indirect mechanism involves SARS-CoV-2 infection leading to the release of DAMPs such as HMGB1(**Figure 1**), which in turn bind to and activate TLR4. More directly, the SARS-CoV-2 spike protein itself functions as a PAMP that activates TLR4 directly, with the binding affinity typical of other direct virus-receptor interactions [14, 15, 34]; moreover, this interaction can be blocked by a small molecule inhibitor of TLR4 [14]. Here we show that both HMGB1 and SARS-CoV-2 Spike protein-mediated TLR4 activation can be inhibited by paridiprubart, reducing the cellular response to near the unstimulated control (**Figure 2b**).

**Figure 2.**
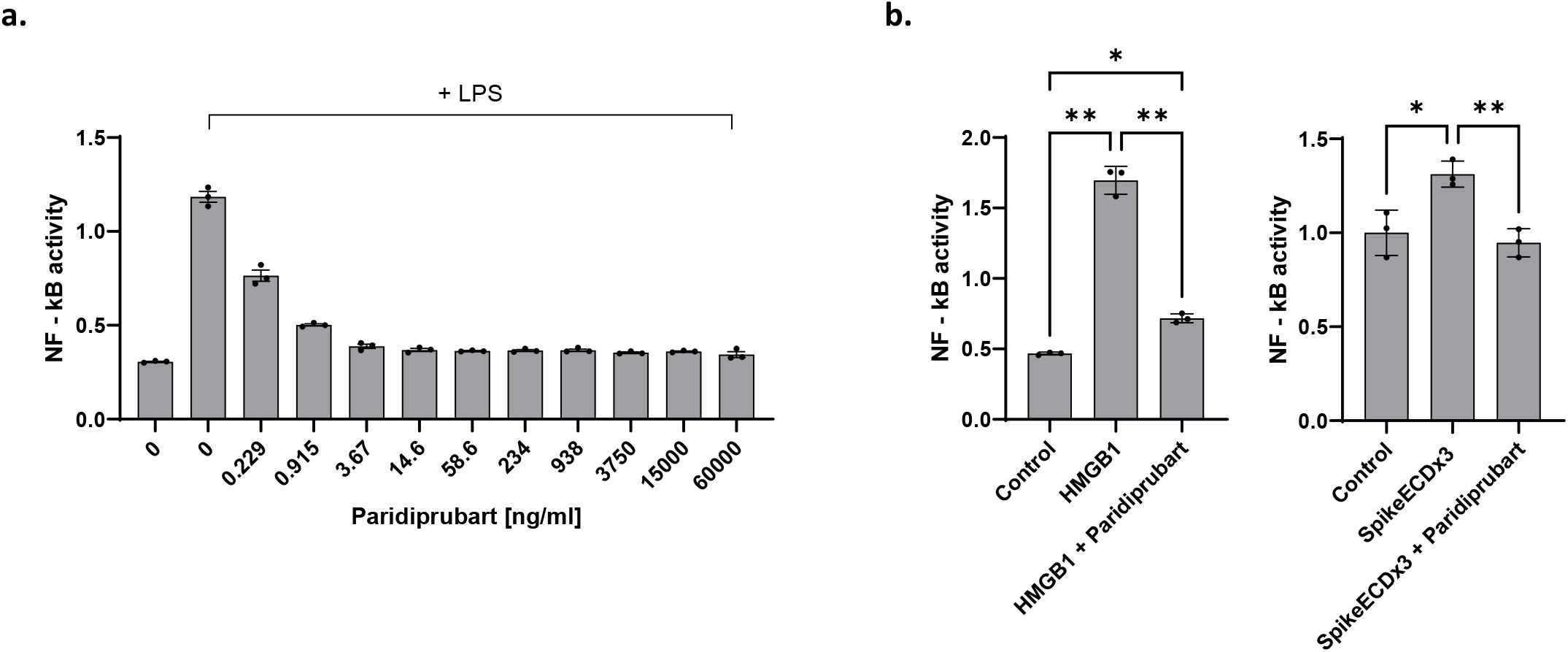
Cellular NF-kB response to TLR4 agonists is blocked by paridiprubart. **A)** TLR4-specific monoclonal antibody paridiprubart blocks NF-kB reporter activation upon exposure to LPS in a dose-dependent manner. **B)** Paridiprubart inhibits HMGB1 (left panel) and SARS-CoV-2 (right panel) activation of NF-κB reporter in THP1 cells. * p<0.05, **p<0.01

### Paridiprubart blocks activation of TLR4 by different pathogens

Taken together, the findings above are consistent with SARS-CoV-2 being able to activate TLR4 signaling directly (by Spike protein) and indirectly (by increasing levels of TLR4 DAMPs). We next tested whether the inflammatory response elicited by SARS-CoV-2 infection, which has the potential to stimulate multiple pattern recognition receptors [43], could be suppressed by paridiprubart. SARS-CoV-2 elicits an NF-κB-driven response in THP1 cells, and this is effectively blocked by paridiprubart treatment, reducing the reporter expression to the background level **(Figure 3a)**.

**Figure 3:**
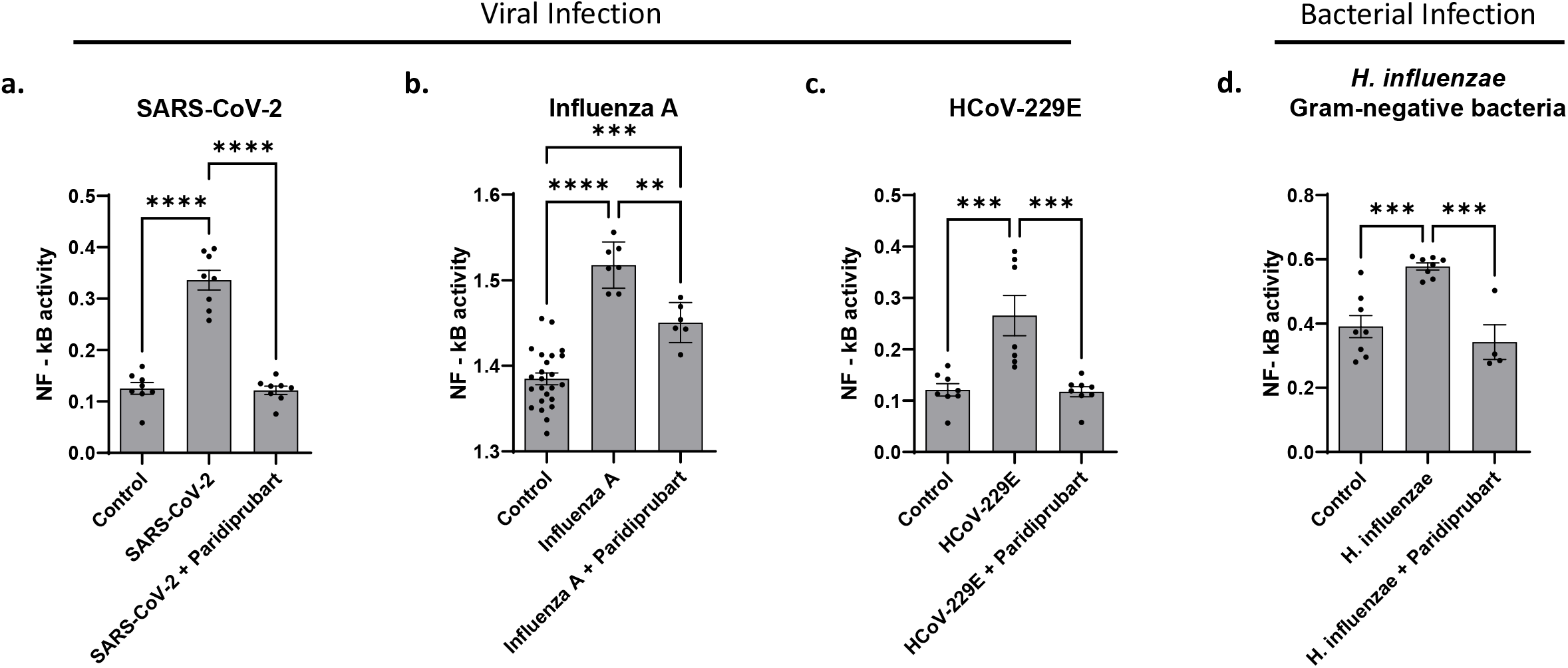
Pathogen-induced NF-kB response is blocked by paridiprubart. Paridiprubart inhibited NF-kB activation elicited by **A)** SARS-CoV-2, **B)** Influenza A virus, **C)** HCoV-229E, or **D)** *Haemophilus influenzae* bacteria in PMA-differentiated THP1 monocytes ** p<0.01, *** p<0.001, **** p<0.0001

Since other pathogens can also activate NF-κB signaling through the stimulation of multiple pattern recognition receptors by their own expression of PAMPs and/or infection-induced DAMPs [10, 18, 44], we considered whether the effect of paridiprubart was specific to SARS-CoV-2 or could also influence the NF-κB response caused by other pathogens. For this purpose, we performed infections with Influenza A (H1N1) virus, the human coronavirus 229E (HCoV-229E), and the gram-negative bacteria *H. influenzae*. In each case, paridiprubart significantly inhibited the NF-κB response, typically causing the treated samples to be indistinguishable from the uninfected controls **(Figure 3b-d)**. Combined, these results indicate that paridiprubart has the capacity to suppress inflammatory response to a variety of pathogens despite their expression of diverse innate immune-stimulatory factors.

## Discussion

Clinical data has revealed that ARDS occurs in SARS patients despite a diminishing viral load, suggesting that antiviral therapy alone may be inadequate to prevent disease progression and even death caused by the aberrant host immune response [45]. Evidence gathered during the COVID-19 pandemic suggests two predominant drivers of COVID-19 pathogenesis. During the early stages of the disease, SARS-CoV-2 replication and viral load are the main driver of disease symptoms. However, as the disease progresses, an overactive, uncontrolled immune response to the initial viral infection can cause a pathogenic inflammation that leads to tissue damage, and which can progress to ARDS and multiple organ failure. Ultimately this cascade results in hospitalization and mortality in a subset of the population [46]. This host-driven inflammatory response has been linked to other severe viral pneumonias that lead to pathophysiological organ failure, including avian influenza, seasonal influenza and severe acute respiratory syndrome (SARS) [4-6]. Overactive inflammatory processes have also been implicated in infectious diseases such as Ebola, Dengue and RSV [10].

While TLR4 was originally considered to be most important for innate detection of bacterial infections, it is now recognized to fulfill a broader function of recognizing both pathogen-derived and danger signals. Since the emergence of SARS-CoV-2, TLR4 has also been shown to play a prominent role in COVID-19 pathogenesis [14, 47]. Following viral-mediated cell damage in the lungs, TLR4-activating DAMPS are released, including calprotectin [31, 48-51], HMGB1 [42], resistin [52] and Ox-PL [18]. These DAMPs are potent activators of TLR4 found on macrophages, neutrophils, dendritic cells and other cells, and their release can lead to a runaway proinflammatory cascade [18]. In addition to SARS-CoV-2 spike protein, the RSV fusion protein, Ebola virus glycoprotein, and dengue virus nonstructural protein 1 are viral glycoproteins that have been observed to activate TLR4 during infection, which may contribute to immunopathogenesis [10, 36]. The combination of direct and indirect triggering of TLR4 during infection presumably explains why the TLR4-specific paridiprubart, which inhibits TLR4 signaling rather than ligand binding [9], so effectively suppressed the inflammatory signals that were otherwise apparent in response to lipopolysaccharide, bacterial infection or viral infection, as shown herein. This raises the hope that paridiprubart may be an effective therapeutic with application against the disease caused by a variety of pathogens.

## Author contribution

Conceptualization R.M., B.G., M.B., I.T., S.D.G.; Methodology R.M.,

R.H., S.A., J.L., S.M., Analysis/Interpretation of Data R.M., R.H., N.R., B.G., S.D.G.; Wrote the manuscript R.M., L.K., J.L., K.K., L.C., N.R., B.G., M.B., S.D.G. All authors contributed and approved the final version of the manuscript.

## Acknowledgement

We would like to thank Drs. Natasha Christie-Holmes, Epshita Islam and

Azmiri Sultana for technical support and advice during the course of this study.

